# A transcriptome-wide association study identifies *PALMD* as a susceptibility gene for calcific aortic valve stenosis

**DOI:** 10.1101/184945

**Authors:** Sébastien Thériault, Nathalie Gaudreault, Maxime Lamontagne, David Messika-Zeitoun, Marie-Annick Clavel, Romain Capoulade, François Dagenais, Philippe Pibarot, Patrick Mathieu, Yohan Bossé

## Abstract

Calcific aortic valve stenosis (CAVS) is a common and life-threatening heart disease with no drug that can stop or delay its progression. Elucidating the genetic factors underpinning CAVS is an urgent priority to find new therapeutic targets^1^. Major landmarks in genetics of CAVS include the discoveries of *NOTCH1*^2^ and *LPA*^3^. However, genetic variants in these genes accounted for a small number of cases and low population-attributable risk. Here we mapped a new susceptibility locus for CAVS on chromosome 1p21.2 and identified *PALMD* (palmdelphin) as the causal gene. *PALMD* was revealed using a transcriptome-wide association study (TWAS)^4^, which combines a genome-wide association study (GWAS) of 1,009 cases and 1,017 ethnically-matched controls with the first large-scale expression quantitative trait loci (eQTL) mapping study on human aortic valve tissues (n=233). The CAVS risk alleles and increasing disease severity were both associated with lowered mRNA expression levels of *PALMD* in valve tissues. The top variant explained up to 12.5% of the population-attributable risk and showed similar effect and strong association with CAVS (P=1.53 × 10^−10^) in UK Biobank comparing 1,391 cases and 352,195 controls. The identification of *PALMD* as a susceptibility gene for CAVS provides new insights about the genetic nature of this disease and opens new avenues to investigate its etiology and develop much-needed therapeutic options.

---

Calcific aortic valve stenosis (CAVS) is the most common valvular heart disease (2% >65 years old)^5^. It is characterized by a progressive remodeling and calcification of the aortic valve, leading to stenosis and heart failure. There is a long latent period before CAVS becomes severe and symptomatic, which provides a window of opportunity for intervention^6^. Unfortunately, conventional cardiovascular drugs are unable to stop or delay the progression of CAVS^7^. If severe CAVS is left untreated, the survival of affected patients is dramatically shortened following the onset of symptoms^6^. The only treatments available for symptomatic patients with severe CAVS are invasive and costly surgical or transcatheter interventions^8^. Finding new molecular targets to halt disease progression is an urgent priority^1^.

Our molecular and genetic understanding of CAVS is currently limited. A strong genetic component is suggested by early-onset forms of the disease, familial aggregation and genetic epidemiology studies^9,10^. A number of susceptibility genes were identified^3,11–14^. Genome-wide gene expression of normal and stenotic human valves have highlighted biological processes that are involved in CAVS^15,16^. In this study, we combined GWAS and valve eQTL results to identify the molecular drivers of CAVS.

Table 1 shows the clinical characteristics of 1,009 cases and 1,017 controls that passed quality controls (QC) in the QUEBEC-CAVS cohort. Mean age was 71.7±8.3 years and 64% were men. About half (53%) of CAVS cases and the majority of controls (98%) had coronary artery disease. The GWAS evaluated 7,732,680 SNPs after imputation. The quantile-quantile plot indicated no inflation of observed test statistics (**Supplemental Fig. 1**). GWAS results are illustrated in Fig. 1a. No SNP reached genome-wide significance (P_GWAS_=5 × 10^−8^). Eight loci had at least one SNP with P_GWAS_<1 × 10^−6^ (**Supplemental Table 1 and Supplemental Fig. 2**).

**Figure 1.**
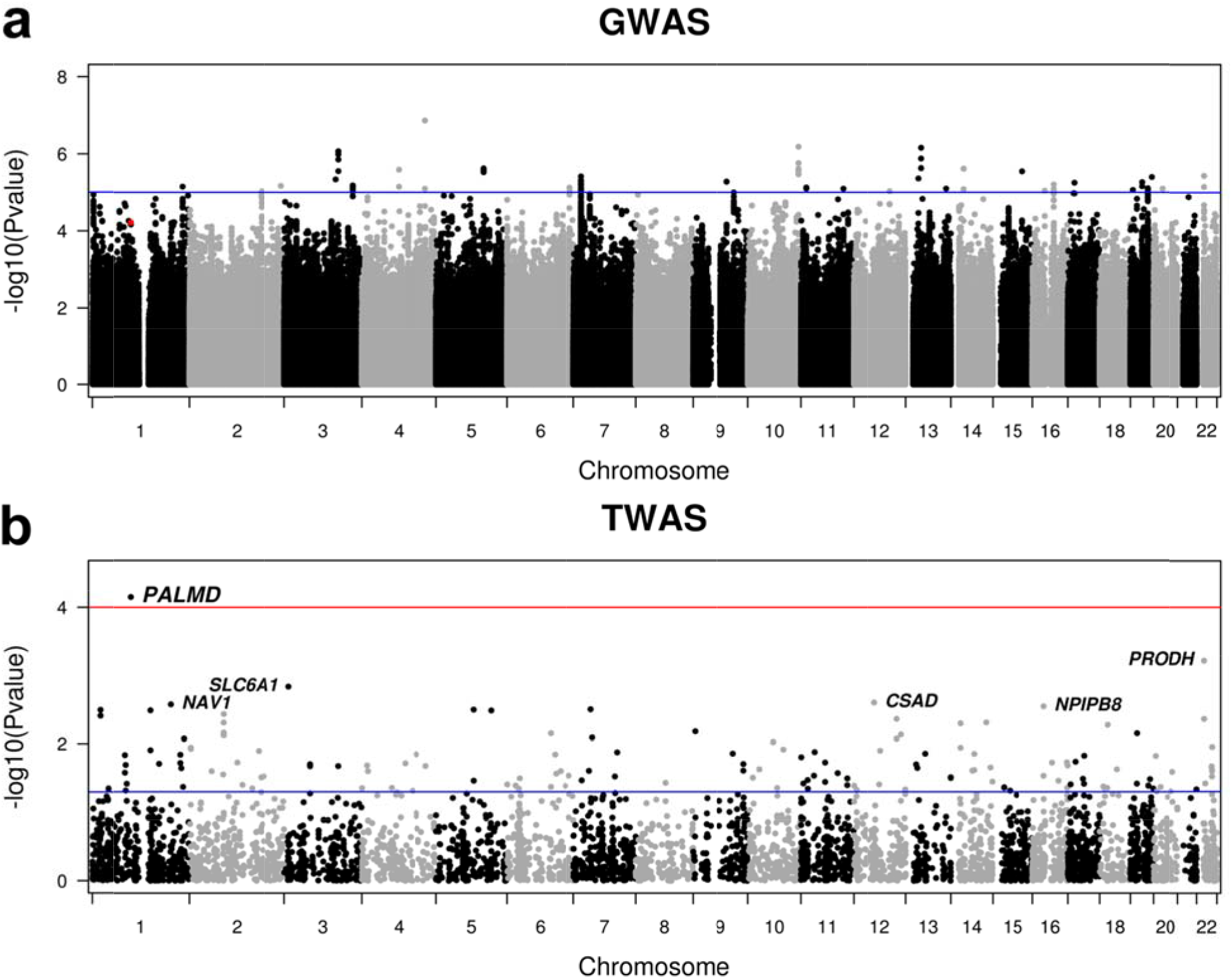
Manhattan plots showing the GWAS in QUEBEC-CAVS and TWAS results. (a) Genetic associations with CAVS observed in 1,009 cases and 1,017 controls. The y axis represents *P* value in -log 10 scale. The horizontal blue line indicates a *P* value of 1 × 10^−5^. The red dot indicates rs6702619. (b) Transcriptome-wide association in valve tissue with CAVS. *P* values for gene expression-CAVS associations are on the y-axis in -log10 scale. The blue horizontal line represents *P_TWAS_* of 0.05. The red horizontal line represents the genome-wide significant threshold used in this study (P_TWAS_<0.0001). Annotations for the top significant probes are indicated.

**Table 1.** Clinical characteristics of the QUEBEC-CAVS cohort

| **Characteristics** | **Cases**<br>**(n = 1,009)** | **Controls**<br>**(n = 1,017)** |
| --- | --- | --- |
| Age (years) | 72.5 ± 8.4 | 70.9 ± 8.1 |
| Gender (% male) | 63.6% | 64.7% |
| Aortic valve area (cm<sup>2</sup>) | 0.77 ± 0.30 (19) | NA |
| Mean gradient (mm Hg) | 41.1 ± 16.0 (64) | NA |
| Concomitant coronary artery bypass grafting (%) | 53.3% | 98.3% |
| Diabetes (%) | 32.4% | 32.4% |
| Hypertension (%) | 75.6% | 75.0% |
| BMI (kg/m<sup>2</sup>) | 28.6 ± 5.4 | 27.7 ± 4.7 |
| Cholesterol (mmol/L) | 4.2 ± 1.1 (30) | 3.9 ± 1.0 (23) |
| Triglycerides (mmol/L) | 1.5 ± 0.9 (32) | 1.5 ± 0.8 (24) |
| LDL-C (mmol/L) | 2.2 ± 0.9 (36) | 2.0 ± 0.8 (31) |
| HDL-C (mmol/L) | 1.3 ± 0.4 (32) | 1.1 ± 0.3 (28) |
| eGFR (mL/min/1.73m<sup>2</sup>) | 66.9 ± 18.2 (1) | 68.6 ± 16.7 (3) |

An expression quantitative trait loci (eQTL) mapping study was then conducted in the most disease-relevant tissue to study CAVS, i.e. human aortic valve. Valve eQTL were calculated in 233 patients that passed QC for genotyping and gene expression. A total of 10,598 independent valve eQTL were identified at *P*_eQTL_<1 × 10^−8^ (**Supplemental Table 2**). Overall, these eQTL involved 2,277 genes/probes significantly associated with one to 41 independent SNPs (r^2^<0.8) (**Supplemental Fig. 3a**). On average, a single SNP explained 26.5% of the gene/probe expression variance. However, 29.1% of the eQTL-SNPs explained more than 30% of the expression variance (**Supplemental Fig. 3b**).

A transcriptome-wide association study (TWAS)^4^ was then performed using the GWAS and valve eQTL datasets. A Manhattan plot showing transcriptome-wide association in valve tissue with CAVS is shown in Fig. 1b. Only one probe-CAVS association corresponding to the *PALMD* gene reached genome-wide significance (P_TWAS_=0.00007). *PALMD* is located on chromosome 1p21.2 and the top GWAS SNPs were the same as the top valve eQTL-SNPs (Fig. 2a). rs6702619, a genotyped variant, was the SNP in this locus most significantly associated with both CAVS (OR=1.29, 95% CI 1.14-1.46, *P*_GWAS_=6.12 × 10^−5^) and the expression of *PALMD* (*P*_eQTL_=5.82 × 10^−33^). rs6702619 is a common SNP (minor allele frequency of 47.7% in Europeans from the 1000 Genomes Project) located 65.2 kb from the transcriptional start site of *PALMD.* The valve eQTL rs6702619-PALMD indicated that the risk allele for CAVS (“G”) is associated with lower mRNA expression levels of *PALMD* in valve tissues (Fig. 2b), suggesting that lower expression increases the risk of CAVS. Variants located within 1 Mb of *PALMD* that increased its expression tended to decrease the risk of CAVS (Fig. 2c). A Mendelian randomization analysis suggested a causal role of lower *PALMD* expression on CAVS risk, without evidence of pleiotropy (P=0.00046; Egger intercept P=0.19; Supplemental **Fig. 4**). Results remained significant after excluding rs6702619 from the analysis (P<0.05). Together, these results indicated that risk variants on 1p21.2 confer susceptibility to CAVS through down-regulation of *PALMD* in aortic valve tissues. The relationship between *PALMD* expression levels and CAVS severity was also consistent with this direction of effect. Lower normalized, age and sex adjusted *PALMD* expression was associated with smaller aortic valve area (P=0.0027), higher mean transvalvular gradient (P=0.0001), and higher peak transvalvular gradient (*P*=8.13 × 10^−5^) (**Supplemental Fig. 5**).

**Figure 2.**
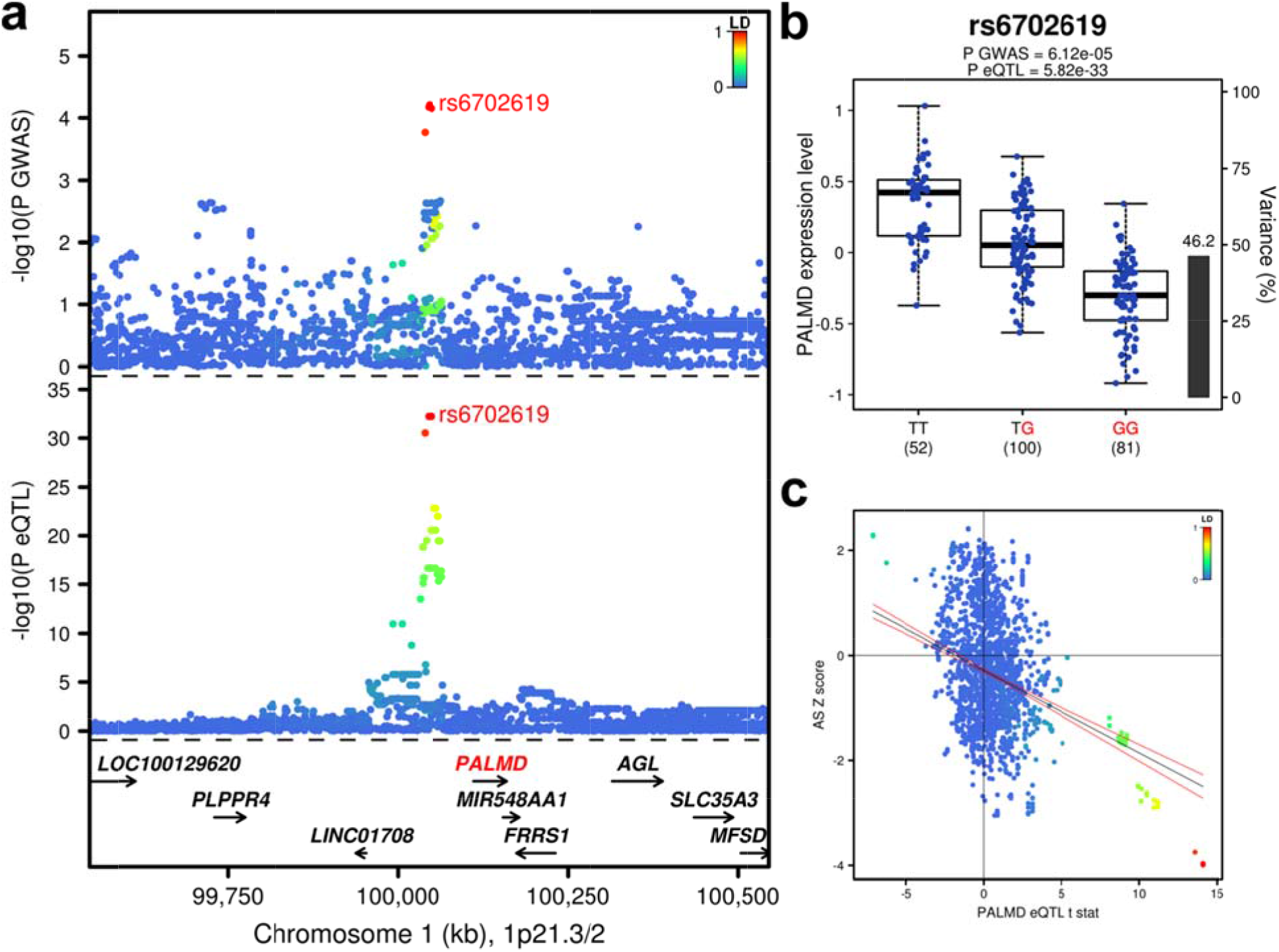
*PALMD* is the causal gene on the 1p21.2 CAVS susceptibility locus. (a) GWAS and valve eQTL results surrounding *PALMD* on chromosome 1p21. The upper panel shows the genetic associations with CAVS. The bottom panel shows the valve eQTL statistics for the *PALMD* gene. The extent of linkage disequilibrium (LD; r^2^ values) for all SNPs with rs6702619 is indicated by colors. The location of genes in this locus is illustrated at the bottom. (b) Boxplots of gene expression levels in the valves according to the three genotype groups for SNP rs6702619. The y-axis shows the mRNA expression levels. The x-axis represents the three genotype groups for rs6702619 with the number of individuals in parenthesis. The risk allele identified in our GWAS is shown in red. Box boundaries, whiskers and center marks in boxplots represent the first and third quartiles, the most extreme data point which is no more than 1.5 times the interquartile range, and the median, respectively. The black bar and the left y axis indicate the variance in *PALMD* gene expression explained by rs6702619. (c) Scatterplot of the 1p21.2 susceptibility locus showing SNP associations with CAVS and *PALMD* gene expression in aortic valve tissues. The y axis represents variant association with CAVS (Z score). The x axis shows association with *PALMD* gene expression (t statistic). Variants are colored based on the degree of LD (r) with the top CAVS-associated variant rs6702619. The blue line is the regression slope with 95% confidence interval (red lines).

We additionally looked for *PALMD* eQTL in the Genotype-Tissue Expression (GTEx) dataset^17^. rs6702619-PALMD association was not significant in 44 tissues from GTEx, suggesting that the change in expression is specific to aortic valve tissue. Evaluating other SNPs within or near *PALMD* revealed significant eQTL with this gene in the pancreas, tibial nerve and subcutaneous adipose tissue. However, these eQTL-SNPs were not in LD with CAVS-associated variants.

We then replicated our newly associated locus using data recently released by the UK Biobank, which included 1,391 CAVS cases and 352,195 unaffected individuals all of European ancestry. We observed a very strong association with an almost identical effect on CAVS risk for our top SNP rs6702619 with an OR of 1.27 (95% CI 1.18-1.37) (Fig. 3a). The association was also replicated for rs7543130, a genotyped variant in perfect LD with rs6702619. At the genome-wide scale level, only two loci were genome-wide significant in UK Biobank (Fig. 3b): *LPA* on 6q25.3-q26 previously associated with aortic valve calcification^3^ and *PALMD* on 1p21.2 identified in this study.

**Figure 3.**
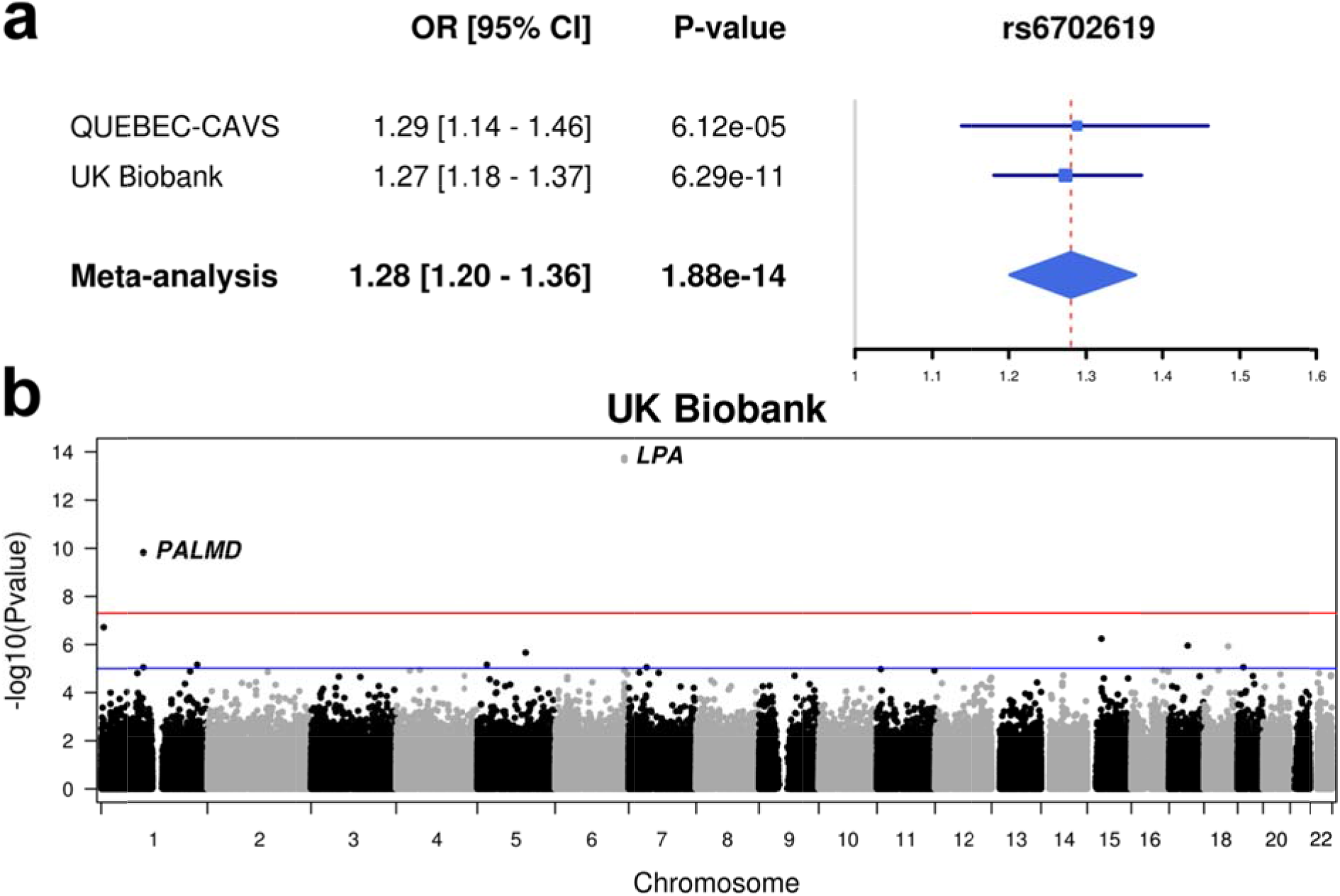
Replication in UK Biobank. (a) Forest plot of overall effect size for rs6702619 in the discovery (QUEBEC-CAVS) and validation (UK Biobank) cohorts. The blue filled squares represent the odds ratio (OR) for each cohort. The horizontal lines represent the 95% confidence intervals of the OR. The grey and the dashed red vertical lines represent an OR of 1.0 and the OR of the meta-analysis, respectively. (b) Manhattan plot showing the GWAS in UK Biobank comparing 1,391 CAVS cases and 352,195 controls. The horizontal blue and red lines indicate *P_GWAS_* values of 1 × 10 and 5 × 10, respectively. Three SNPs were significant at the *P_GWAS_* threshold of 5 × 10^−8^: rs74617384 and rs10455872 at the *LPA* locus and rs7543130 (in perfect LD with rs6702619) at the *PALMD* locus.

The top GWAS SNP in this study (rs6702619) was recently identified by the EchoGen consortium to be associated with aortic root diameter in a large GWAS meta-analysis including up to 46,533 individuals^18^. The EchoGen consortium also identified a SNP in *CACNA1C* (calcium channel, voltage-dependent, L type, alpha 1C subunit) associated with aortic root diameter^18^, which we recently identified as a susceptibility gene of CAVS^14^. As indicated in the later study, rs6702619 is located within enhancer histone marks and DNase-hypersensitive sites in Encyclopedia of DNA Elements (ENCODE) data^19^. However, rs6702619 was not found to act as *cis*-eQTL in whole blood, monocytes, and myocardial tissue^18^. In this study, we found *PALMD* consistently expressed in human aortic valve tissues and rs6702619 strongly associated with mRNA expression levels of *PALMD*, providing a potential functional mechanism of how genetic variants on 1p21.2 may increase CAVS susceptibility. Furthermore, a Mendelian randomization analysis suggested that *PALMD* expression is causally associated with the risk of CAVS.

The frequency of rs6702619 is high in our French Canadian population (MAF=47%) and the modest effect size (OR=1.29) is consistent with those observed for other complex traits. The estimated population-attributable risk indicates that more than 12.5% of cases in our population and 12.3% of cases in UK Biobank could be attributed to this common variant. Of note, the frequency of rs6702619 varies greatly between ethnic groups (**Supplemental Fig. 6**). In the 1000 Genomes Project, allele “G” had a frequency of 48% in Europeans, 8% in Africans, 7% in East Asians and 25% in South Asians. This may explain some of the discrepancy in risk of CAVS among ethnic groups^20,21^.

*PALMD* is a distant homolog of a small paralemmin protein family^22^. *PALMD* is expressed in many tissues, but most abundantly in cardiac and skeletal muscle^23^. It is a mainly cytosolic protein, localized predominantly in actin filaments, and may be implicated in plasma membrane dynamics and cell shape control as other members of this family^22,24^. However, its molecular and cellular functions remain largely unknown. *PALMD* was recently shown to promote myoblast differentiation and muscle regeneration^25^. Considering the common embryologic origin of the aortic valve and the aortic root, which both arise from the secondary heart field, a potential role on smooth muscle and valve interstitial cell differentiation could explain the association with the two phenotypes^26,27^. *PALMD* was also identified as a pro-apoptotic gene induced by p53 in response to DNA damage in osteosarcoma cell lines^28^. However, its role in cell death was not confirmed in other cell types^25^.

Of note, the identified SNP, rs6702619, was not associated with aortic valve calcification measured by computed tomographic scanning in a large GWAS including 6,942 individuals of European ancestry (P=0.557)^3^. This suggests that calcification is not the main mechanism by which *PALMD* expression modulates CAVS progression and could rather appear only in later stages of the disease. The variant identified is also expected to act via pathways not involved in the risk of coronary artery disease (CAD) since the control individuals included in our study almost all had CAD. rs6702619 was indeed not associated with CAD in a recent large GWAS meta-analysis (P=0.96)^29^.

Our study has some limitations. First, only individuals of European ancestry were included in the analysis, therefore results cannot be generalized to other ethnic groups. Second, the power in our GWAS analysis was limited by the number of available samples. However, expression data on the most disease-relevant tissue and replication in a large population-based cohort confirmed the robustness of the findings. Third, aortic valve tissue was only available in stenotic valves; patterns of expression of *PALMD* in normal valves might differ.

In conclusion, using a TWAS approach with expression data in aortic valve tissue, we identified a new CAVS susceptibility gene, *PALMD*, with replication in a large prospective population-based cohort. Further analyses of gene expression in valve tissue and disease severity suggested that the identified variant acts by decreasing the expression of *PALMD*, a finding supported by Mendelian randomization. Further studies are warranted to elucidate the exact mechanism of action and evaluate the potential for targeted therapeutic interventions.

## Acknowledgements

We thank the research team at the cardiac surgical database and biobank of the Institut universitaire de cardiologie et de pneumologie de Québec (IUCPQ) for their valuable assistance. This research has been conducted using the UK Biobank Resource. P.P. holds the Canada Research Chair in Valvular Heart Disease and his research program is supported by a Foundation Scheme Grant from Canadian Institutes of Health Research (Ottawa, Ontario, Canada). P.M. holds a Fonds de Recherche du Québec-Santé (FRQS) Research Chair on the Pathobiology of Calcific Aortic Valve Disease. Y.B. holds a Canada Research Chair in Genomics of Heart and Lung Diseases.

## Author contributions

S.T., D.M.Z., P.P., P.M. and Y.B. contributed to the conception and study design.

N.G., D.M.Z., M.A.C., R.C., F.D., P.P., P.M. and Y.B. contributed to data collection.

S.T., M.L. and Y.B. contributed to data analysis.

S.T., N.G., M.L., P.M. and Y.B. contributed to data interpretation.

## Methods

### Study cohort

Blood samples and aortic valves were collected from patients with severe aortic valve stenosis undergoing aortic valve replacement at the *Institut universitaire de cardiologie et de pneumologie de Québec* (QUEBEC-CAVS). Only cases with tricuspid nonrheumatic CAVS were included. No severe regurgitation or other severe valvular heart diseases were present. In parallel, an ethnically-matched control group was recruited from patients that underwent cardiac surgery, mostly for isolated coronary artery bypass (>98%). Other indications for surgery in the control group included heart transplant, tumor removal, aortic endoprosthesis and interatrial communication. Absence of CAVS was confirmed by echocardiography. This cohort of control patients was also matched in a 1:1 ratio with cases for age, gender, type 2 diabetes and hypertension. Patients with a history of severe valvular heart disease (at any of the 4 valves), with significant aortic valve regurgitation (grade > 2) or with end stage renal disease (eGFR<15 mL/min/1.73m^2^) were excluded. All patients signed an informed consent for the realization of genetic studies. Demographics, anthropometric measurements, lifestyle factors, previous and current medical history, current medication and blood pressure measurements were collected. In addition, plasma lipids and creatinine were measured. CAVS cases underwent a comprehensive Doppler echocardiographic examination. The transvalvular gradient was calculated using the modified Bernoulli equation and the aortic valve area calculated with the continuity equation. For this study, 1,033 cases and 1,037 controls were available.

### Genome-wide association study

Blood samples were collected and DNA was extracted from frozen buffy coat. Whole-genome genotyping was performed using the Illumina HumanOmniExpress BeadChip. Standard genotyping quality controls were performed. SNP genotyping data were filtered for call rate <97%, low-quality loci with 10^th^ percentile of Illumina GenCall score ≤0.1, Hardy-Weinberg equilibrium P<1 × 10^−7^, minor allele frequency (MAF) <1% and different call rate between cases and controls (P<1 × 10^−6^). A total of 613,862 SNPs passed quality control checks. Samples were excluded after consideration for the 10^th^ percentile of Illumina GenCall score ≤0.2, genotype completion rate <95%, outlier heterozygosity rate (F>0.20), genotypic and phenotypic gender mismatch, genetic background outliers detected by principal component analysis with HapMap subjects as population reference panel, unexpected duplicates and genetic relatedness (first-degree relatives) evaluated by identity-by-state using PLINK^31^. After the quality control filters, 1,009 cases and 1,017 controls were available for subsequent analyses. A summary of genotyping quality controls on SNPs and samples is provided in **Supplemental Tables 3** and **4**, respectively. Genotypes were then imputed with the Michigan Imputation Server^32^ using the Haplotype Reference Consortium version 1 (HRC.r1-1) data^33^ as reference set (2016-03-31). Variants with an r^2^ value of ≤0.3 or MAF <1% were removed from further analysis. A total of 7,732,680 imputed SNPs passed quality control. Genetic association tests were performed using additive logistic regression models based on expected genotype counts (dosages) as implemented in the software SNPTEST v2.5.2^34^, adjusting for age, sex, and the first ten ancestry-based principal components. The genomic inflation factor for the main case-control analysis was 1.03. The genome-wide significant *P* value cutoff was set to 5 × 10^−8^. Regional plots were created with LocusZoom^35^. The population-attributable risk was calculated as PAR% = 100% × P × (OR - 1)/[P × (OR - 1) + 1], where P is the frequency of the risk allele associated with CAVS in the control group, and OR is the odds ratio calculated in the case-control cohort.

### Gene expression analysis in human aortic valves

Transcriptomic analyses were performed from 240 stenotic aortic valves collected as mentioned above. All stenotic valves were tricuspid and had a fibro-calcific remodeling score of 3 or 4^36^. Selected valves were from cases that are part of the GWAS and included 120 men and 120 women. RNA was extracted from valve leaflets, and gene expression evaluated using the Illumina HumanHT-12 v4 Expression BeadChip. Standard microarray processing and quality control analysis was performed^37^. The raw data were quantile normalized after log2-transformation with the lumi package in R^38^. Only one sample failed quality control, leaving 239 samples for subsequent analyses. Probe sequences were mapped to RefSeq B38, GENCODE v24 B38, mRNA B38, and the human genome (GRCh B38) using Bowtie, and probes not mapping to any coding region were removed. A total of 45,699 probes remained after this step. Expression data were then adjusted for age and sex using residuals obtained with the robust fitting of linear models function (rlm) in the R statistical package MASS. Residual values deviating from the median by more than 3 SD were filtered out.

### Expression quantitative trait loci

Only subjects that passed genotyping and gene expression quality controls were considered for eQTL analysis leaving a sample size of 233. eQTL were identified by using linear regression model and additive genotype effects as implemented in the Matrix eQTL package in R^39^. *Cis-* eQTL were defined by a 2 Mb window, i.e. 1 Mb distance on either side of the SNP. eQTL were calculated on adjusted expression traits to obtain test statistics, *P* values and false discovery rate. Estimates of effect sizes were obtained with PLINK.

### Transcriptome-wide association study (TWAS)

The TWAS was performed using FUSION^4^. Briefly, the valve eQTL was the reference data including 233 individuals that passed QC for both gene expression and genotyping. This reference set was used to calculate gene expression weights, which were computed one probe at a time using SNP genotyping data located 500 kb on both sides of the probe using prediction models implemented in FUSION including 1) the single most significant valve eQTL-SNP as the predictor (top1), 2) LASSO regression, and 3) elastic net regression (enet). All probes that passed QC in the valve eQTL were evaluated (n=45,699). Expression weights were then combined with summary-level GWAS results to estimate association statistics between gene expression and CAVS. Genome-wide significant TWAS genes were considered at *P_TWAS_*<0.0001.

### Mendelian Randomization

We first selected variants located within 1 Mb of the gene of interest identified in the TWAS, *PALMD*, and significantly associated with *PALMD* gene expression at a threshold of *P*<0.05. We kept only independent variants at a linkage disequilibrium threshold of r^2^ < 0.05. Mendelian randomization was performed using the Wald method by regressing genetic effect estimates for CAVS risk as determined in the GWAS analysis (dependent variable) on genetic effect estimates for *PALMD* gene expression as determined in the eQTL analysis. Effect estimates were adjusted for the minor allele frequency of each variant. To determine the significance of the association, a bootstrap method was used. Predicted effects on CAVS risk and *PALMD* gene expression were sampled from a normal distribution with mean and standard deviation as determined from the GWAS and eQTL analyses. A two-tailed *P* value was calculated using 100,000 random simulations. To determine the presence of unmeasured net pleiotropy, we performed Egger Mendelian randomization in which a non-zero y-intercept is allowed in order to assess violations of standard Mendelian randomization^40^.

### Gene expression according to severity

CAVS disease severity was assessed by aortic valve area, mean and peak transvalvular gradients. The influence of normalized, age and sex adjusted *PALMD* expression levels on disease severity was tested in 239 cases using linear regression models. 95% confidence intervals were estimated using the *predict* function in R.

### Replication in UK Biobank cohort

UK Biobank is a large prospective cohort of about 500,000 individuals between 40 and 69 years old recruited from 2006 to 2010 in several centres located in the United Kingdom^41^. The present analyses were conducted under UK Biobank data application number 25205. We used genotyping data obtained from the second genetic data release including 488,377 individuals. Samples were genotyped with the Affymetrix UK BiLEVE Axiom array or the Affymetrix UK Biobank Axiom Array. Phasing and imputation were performed centrally using a reference panel combining the Haplotype Reference Consortium (HRC) as a first choice and UK10k and 1000 Genomes Phase 3 samples for SNPs not available in HRC. Samples with call rate <95%, outlier heterozygosity rate, gender mismatch, non-white British ancestry, related samples (second degree or closer), samples with excess third degree relatives (>10) or not used for relatedness calculation were excluded. Variants not on both arrays, which failed quality control in more than one batch, with call rate <95% or with minor allele frequency <0.0001 were excluded.

CAVS diagnosis was established from hospital record, using the International Classification of Diseases version-10 (ICD10) and Office of Population Censuses and Surveys Classification of Interventions and Procedures (OPCS-4) coding. ICD10 code number I35.0 and OPCS4 code number K26 were used. Participants with a history of rheumatic fever or rheumatic heart disease as determined by ICD10 codes I00 to I02 and I05 to I09 were excluded.

We performed additive logistic regression analysis using SNPTEST v2.5.2, adjusting for age, sex, and the first ten ancestry-based principal components to evaluate the effect of the top SNP identified in our TWAS analysis. We then performed a fixed-effect meta-analysis using the inverse-variance weighted method. To look for other significant variants associated with CAVS in UK Biobank, we performed a genome-wide association study from the genotyped data using additive logistic regression models in PLINK adjusting for age, sex and the first ten principal components. The genomic inflation factor for the main case-control analysis was 1.01. The genome-wide significant *P* value cutoff was set to 5 × 10^−8^.

### Statistical analysis

Statistical analyses were performed with R version 3.2.3 unless otherwise specified. 2-sided *P* values below 0.05 were considered significant unless otherwise specified.

